# A draft genome of grass pea (*Lathyrus sativus*), a resilient diploid legume

**DOI:** 10.1101/2020.04.24.058164

**Authors:** Peter M. F. Emmrich, Abhimanyu Sarkar, Isaac Njaci, Gemy George Kaithakottil, Noel Ellis, Christopher Moore, Anne Edwards, Darren Heavens, Darren Waite, Jitender Cheema, Martin Trick, Jonathan Moore, Anne Webb, Rosa Caiazzo, Jane Thomas, Janet Higgins, David Swarbreck, Shiv Kumar, Sagadevan Mundree, Matt Loose, Levi Yant, Cathie Martin, Trevor L. Wang

**Affiliations:** John Innes Centre, Norwich Research Park, Colney Lane, Norwich, NR4 7UH, United Kingdom; Biosciences eastern and central Africa International Livestock Research Institute Hub, ILRI campus, Naivasha Road, P.O. 30709, Nairobi 00100, Kenya; Earlham Institute, Norwich Research Park, Colney Lane, Norwich, NR4 7UZ, United Kingdom; University of Nottingham, University Park, Nottingham, NG7 2RD, United Kingdom; National Institute of Agricultural Botany, Huntingdon Road, Cambridge, CB3 0LE, GB; International Center for Agricultural Research in the Dry Areas, Avenue Hafiane Cherkaoui, Rabat, Morocco; Queensland University of Technology, 2 George St, Brisbane City QLD 4000, Australia

**Keywords:** grass pea, *Lathyrus*, legume, beta-ODAP, genome

## Abstract

We have sequenced the genome of grass pea (*Lathyrus sativus*), a resilient diploid (2n=14) legume closely related to pea (*Pisum sativum*). We determined the genome size of the sequenced European accession (LS007) as 6.3 Gbp. We generated two assemblies of this genome, i) EIv1 using Illumina PCR-free paired-end sequencing and assembly followed by long-mate-pair scaffolding and ii) Rbp using Oxford Nanopore Technologies long-read sequencing and assembly followed by polishing with Illumina paired-end data. EIv1 has a total length of 8.12 Gbp (including 1.9 billion Ns) and scaffold N50 59,7 kbp. Annotation has identified 33,819 high confidence genes in the assembly. Rbp has a total length of 6.2 Gbp (with no Ns) and a contig N50 of 155.7 kbp. Gene space assessment using the eukaryote BUSCO database showed completeness scores of 82.8 % and 89.8%, respectively.

## Introduction

Grass pea (*Lathyrus sativus* L.), is a diploid legume (2n=14) first domesticated in the Balkan peninsula (Kislev 1989). It is grown for its seeds as food and feed, as well as for fodder, mainly by farmers in the Indian subcontinent as well as in northern African countries such as Ethiopia (Campbell 1997; Kumar et al. 2011). It is self-fertile, though under field conditions it is often cross pollinated by insects such as bees. Grass pea shows remarkable tolerance to environmental stress, including both drought and waterlogging (Yadav et al. 2006; Campbell 1997) making it a vital source of food and feed in times of scarcity, demonstrated repeatedly during its 8000 years of cultivation (Campbell 1997). It is highly efficient at fixing atmospheric nitrogen in soils using rhizobial symbionts, meaning that it has low input requirements for fertilizers (Jiao et al. 2011; Drouin, Prëvost, and Antoun 2000). For many, grass pea represents an ‘insurance crop’ as it survives and produces a yield under conditions, such as drought or flooding, where most other crops fail (Girma, Tefera, and Dadi 2011; Zhelyazkova et al. 2016; Silvestre et al. 2014; Yang and Zhang 2005; Vaz Patto et al. 2006) and it is often grown by poor farmers using minimal inputs to ensure a supply of food. Its seeds are high in protein (up to 30% w/w in dry seed) (Emmrich 2017), and it has the potential to provide nutritional food security in some of the most resource scarce and least developed regions in the world in the face of climate change and increasing population pressure (Sarkar et al. 2019).

The chief drawback of grass pea as a food has been the presence of an anti-nutritional factor, β-N-oxalyl-L-α,β-diaminopropionic acid (β-ODAP) in most tissues of the plant, including seeds. In conjunction with severe malnutrition, prolonged consumption of β-ODAP causes a neurological disorder, neurolathyrism, which results in permanent paralysis of the lower limbs in humans (Dufour 2011; Kusama-Eguchi et al. 2014). This has led to bans on grass pea cultivation and commerce (Cohn and Streifler 1983), leading to underinvestment in research in this promising legume. The main thrust of grass pea research has been the elimination of β-ODAP from grass pea. A number of low β-ODAP grass pea cultivars have been developed by plant breeders, such as BioL-212 (Ratan), Mahateora, LS 8246, Prateek and Ceora (Kumar et al. 2011; Sawant, Jayade, and Patil 2011; Chakrabarti, Santha, and Mehta 1999; Santha and Mehta 2001; Tsegaye, Tadesse, and Bayable 2005; Siddique, Hanbury, and Sarker 2006), but no zero β-ODAP grass pea cultivar has been produced to date. The biosynthesis of β-ODAP in grass pea is partially understood (Lambein et al. 1993; Kuo and Lambein 1991; Ikegami et al. 1999; Ikegami et al. 1993; Malathi, Padmanab, and Sarma 1970) but critical gaps remain in our knowledge of the genetics of β-ODAP biosynthesis in grass pea.

We present a draft genome sequence of a European accession (LS007) of grass pea which will be valuable for the identification of genes in the β-ODAP biosynthesis pathway, as well as in identification and selection of traits for agronomic improvement. The data will allow comparative genomic analyses between legumes, help in the development of high quality genetic and physical maps for marker-assisted and genomic selection strategies and enable genome editing and TILLING platforms for grass pea improvement. The availability of this draft genome sequence will facilitate research on grass pea with the goal of developing varieties that fulfil its potential as a high protein, low input, resilient, climate smart crop, suitable for small-holder farmers.

## Results

### Genome size estimation

Using *Pisum sativum* as a standard with a genome size of 4.300 Gbp (Leitch et al. 2019), we undertook flow cytometry and estimated the genome size of grass pea genotype LS007 as 6.297 Gbp ± 0.039 Gbp (stdev.). Peak coefficients of variance were below 3% in all three replicates (see Fig. 1).

**Figure 1.**
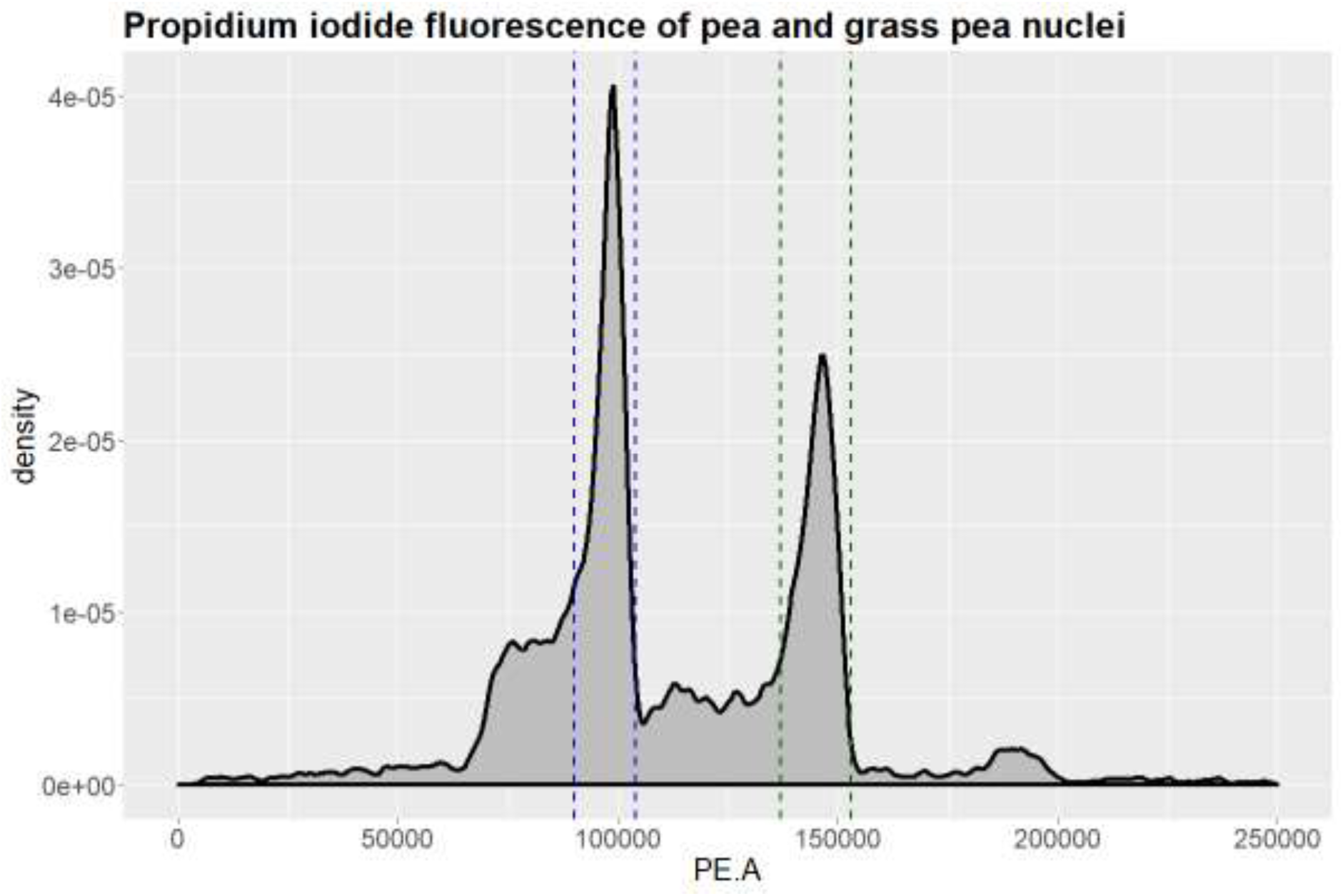
Results of a representative flow cytometry run, after gating to exclude cell debris events. Propidium iodide fluorescence amplitude (in arbitrary units) is shown against event density. The interval assumed to be pea nuclei is defined by blue dashed lines, the interval assumed to be grass pea nuclei is defined by green dashed lines. The experiment was replicated three times.

We used this estimate to calculate genome sequencing depth of all our sequencing datasets. As this estimate is based on *Pisum sativum* as a standard, any inaccuracy in the *Pisum sativum* genome size estimate will affect the genome size estimate for LS007. The Kew Gardens c-value database records 13 *Pisum sativum* samples with haploid genome sizes ranging from 3.724 Gbp to 5.782 Gbp (Leitch et al. 2019). Applied to the grass pea genome this would result in a genome size range of 5.456 Gbp to 8.471 Gbp.

### Sequencing and Genome assembly

Figures 2 and 3 summarise the workflow that generated the datasets and the strategies used to assemble the two versions of the grass pea genome we discuss here, one based entirely on Illumina HiSeq paired-end data scaffolded using Illumina HiSeq long-mate-pair data (EIv1) and one based on Oxford Nanopore Technologies PromethION data assembled de novo, followed by polishing using the Illumina paired-end data.

**Fig. 2.**
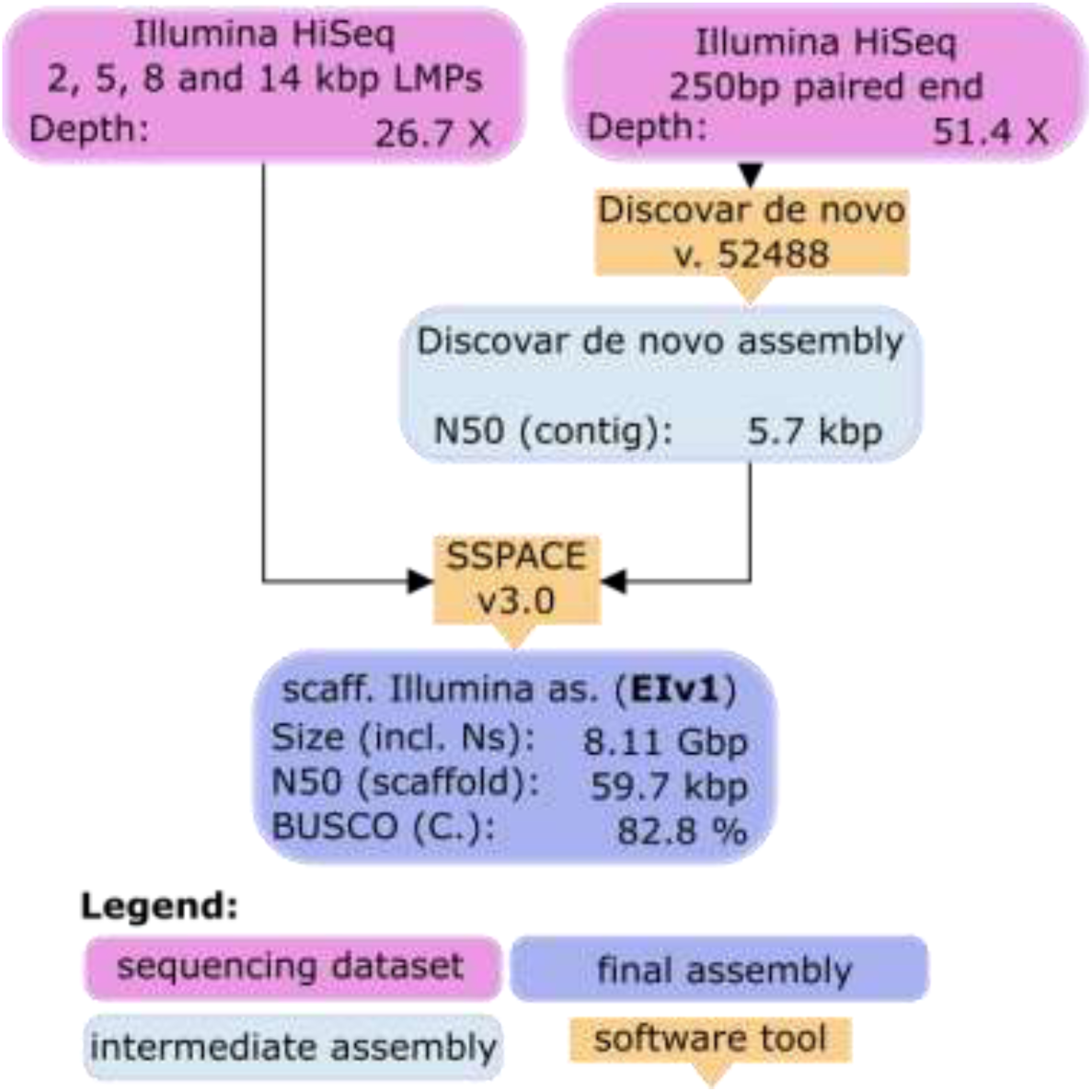
Workflow of assembly and scaffolding of EIv1 from Illumina paired-end and LMP data

**Figure 3.**
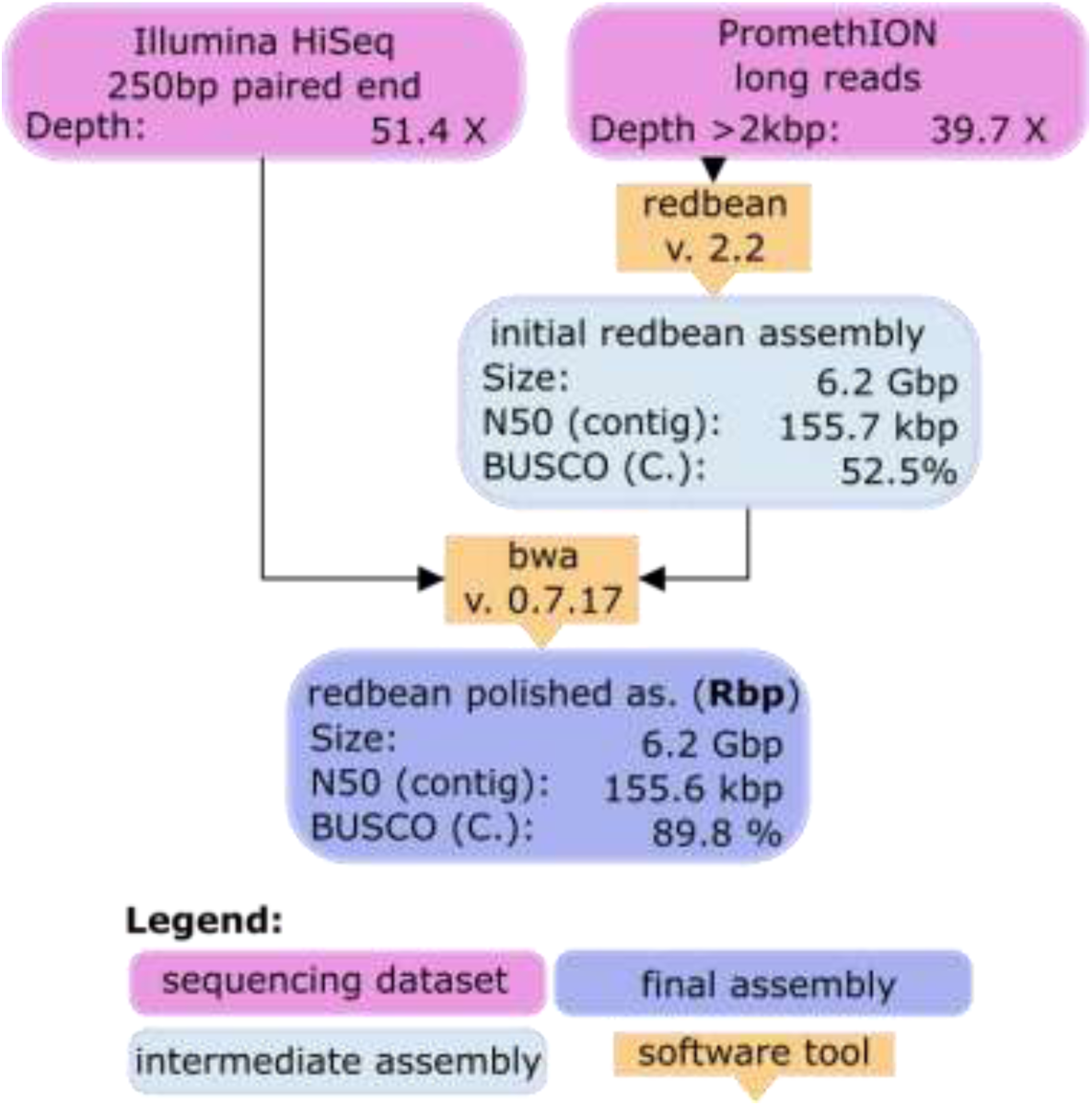
Workflow of assembly of Rbp from ONT data and polishing with Illumina paired-end data.

#### Paired-end sequencing using Illumina platform

The paired end (PE) sequencing of LS007 genomic DNA was carried out using PCR-free libraries, followed by sequencing of the Long Mate Pair libraries (LMP) on the HiSeq Illumina platform (see Material and Methods for details).

#### Assembly

The final EIv1 assembly consisted of 669,893 contigs (minimum size 1 kbp), with a scaffold N50 value of 59,728 bases for a total assembly of 8.12 Gbp, including 1.9 Gbp of Ns in scaffolds.

### Long-read sequencing using the PromethION platform

#### Sequence data

Sequence yields for each load of the flowcells are shown in Table 1. Subsequent loads on the same flowcell were separated by nuclease flushes. In total, 296.15 Gbp of sequence passed the quality filter, representing 47.01 X coverage of a 6.3 Gbp genome (see results section). Distributions of read lengths (post-filter) are shown below.

**Table 1.** ONT PromethION sequencing yield.

| Flowcell/Nuclease<br>flush | DNA preparation | Library<br>amount (ng) | Read N50<br>(kbp) | Yield (Gbp)<br>(passed) |
| --- | --- | --- | --- | --- |
| FC1 1st load | CN-AMP | 250 | 7.84 | 51.84 |
| FC1 2nd load | CN-NS-AMP | 250 | 5.25 | 44.41 |
| FC1 3rd load | QD-SRE | 250 | 18.78 | 14.28 |
| FC2 | QD-SRE | 250 | 22.84 | 57.48 |
| FC3 1st load | CN-NS-AMP-SRE | 270 | 22.31 | 41.7 |
| FC3 2nd load | Mix of CN-NS-AMP-SRE & QD-SRE | 324 | 22.81 | 23.04 |
| FC4 | CN-NS-AMP-SRE | 400 | 29.08 | 32.7 |
| FC5 | CN-NS-AMP-SRE | 400 | 24.71 | 30.7 |

#### Assembly

After filtering for reads >5 kbp, Redbean (Ruan and Li 2020) produced an assembly of 6.2 Gbp, based on 35.8 X coverage. The resulting assembly contained 162,985 contigs, with a contig N50 of 155,574 bp. Following polishing with minimap2 (Li 2018) and bwa (Li 2013), the assembly was brought to a size of 6.237 Gbp in 162,994 contigs with an N50 of 157,998 bp.

### Assembly comparison

Comparison of the EIv1 and the polished Redbean assemblies revealed very similar number of ATGC bases (approx. 6.2 Gbp), the size difference being mostly accounted for by the approximately 1.9 billion Ns in the EIv1 assembly (see Table 2). The GC fraction was 0.388 for the polished Redbean assembly and 0.383 for the EIv1 assembly. Assembly statistics are shown in Figure 4.

**Table 2.**
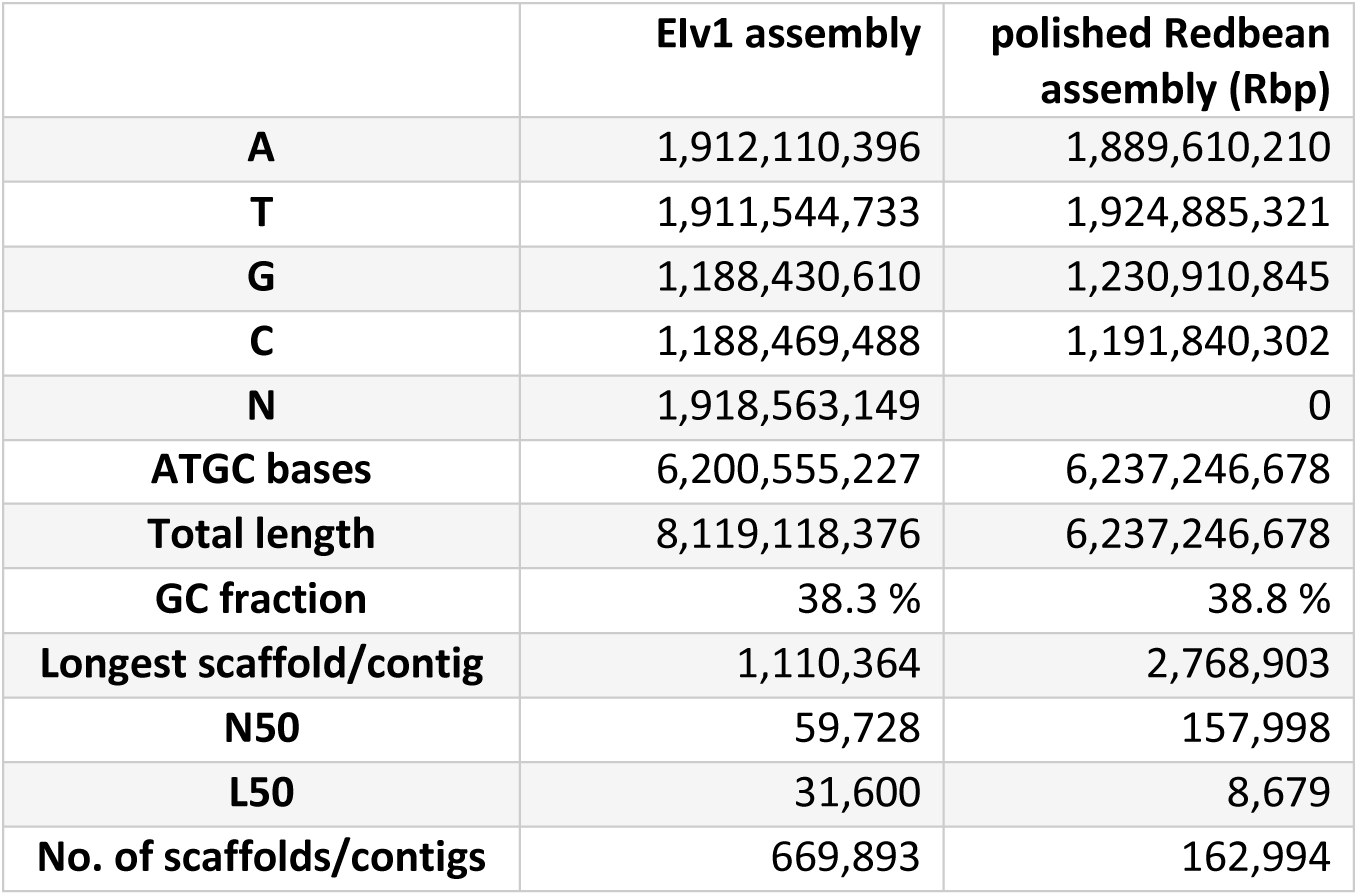
Statistics of the EIv1 and Rbp assemblies.

**Figure 4.**
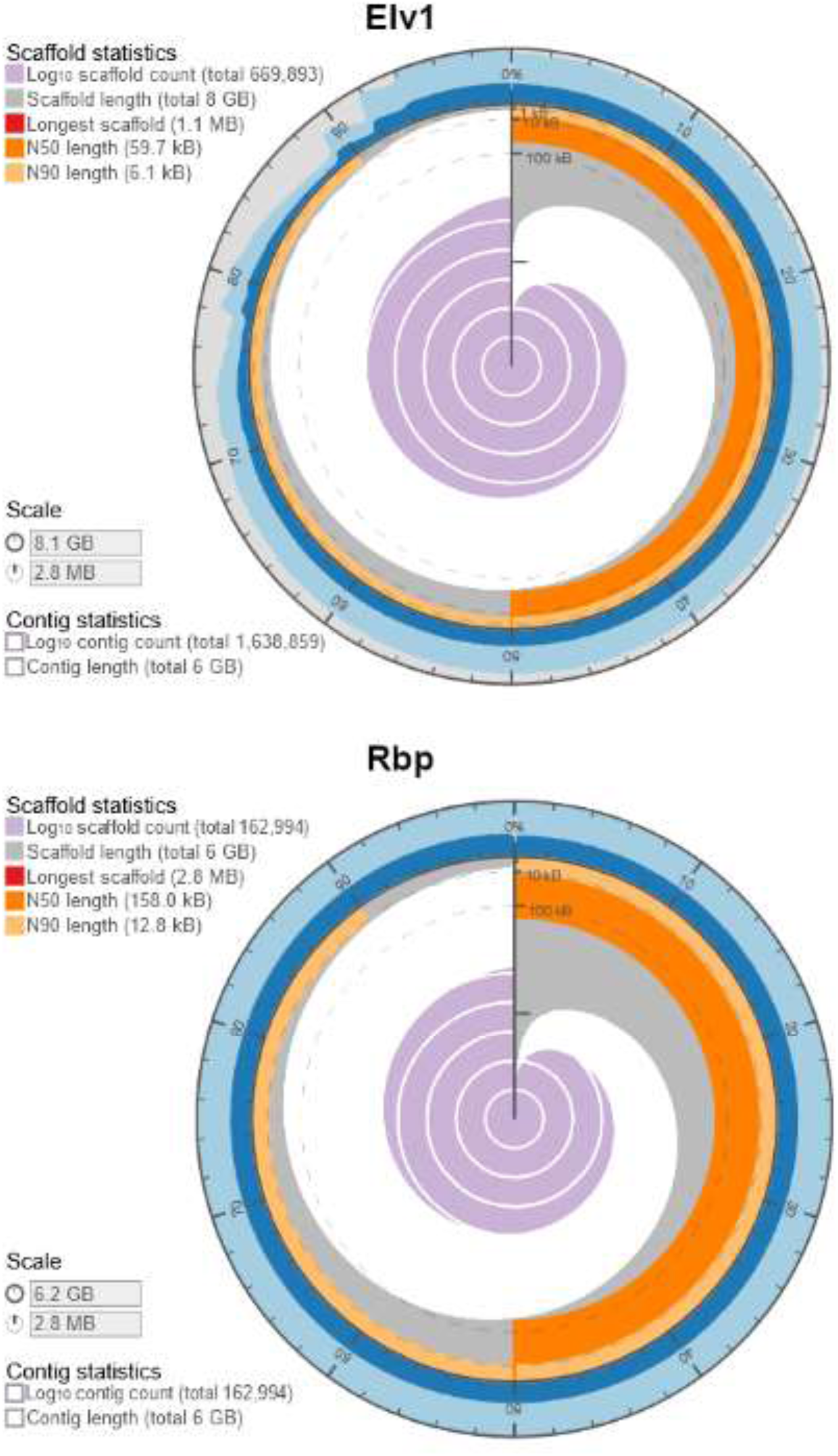
Statistics of the EIv1 and Rbp assemblies of the LS007 genome, visualised using assembly-stats (Challis, 2015). Note the difference in scale for contig/scaffold size.

### BUSCO analysis

BUSCO (Benchmarking Universal Single-Copy Orthologs) analysis was carried out on both the Illumina-only (EIv1) and long-read Redbean assembly polished with Illumina paired-end data (Rbp) to assess the genome assemblies for gene space completeness (Simão et al. 2015). Results are shown in Figure 5 and Table 3.

**Table 3.** BUSCO analysis results using BUSCO:v4.0.4_cv1 and odb10 databases.

| Database | Fabales<br>(n:5366) |  | Eudicots<br>(n:2326) |  | Viridiplantae<br>(n:425) |  | Eukaryote<br>(n:255) |  |
| --- | --- | --- | --- | --- | --- | --- | --- | --- |
|  | Elv1 | Rbp | Elv1 | Rbp | Elv1 | Rbp | Elv1 | Rbp |
| **Complete BUSCOs (C)** | **79.9%** | **82.6%** | **81.7%** | **85.7%** | **86.4%** | **88.4%** | **82.8%** | **89.8%** |
| - Single copy BUSCOs (S) | 66.7% | 80.0% | 68.2% | 82.7% | 71.3% | 84.9% | 66.3% | 82.0% |
| - Duplicated BUSCOs (D) | 13.2% | 2.6% | 13.5% | 3.0% | 15.1% | 3.5% | 16.5% | 7.8% |
| **Fragmented BUSCOs (F)** | **3.9%** | **2.2%** | **6.7%** | **3.0%** | **9.9%** | **3.8%** | **8.2%** | **3.9%** |
| **Missing BUSCOs (M)** | **16.2%** | **15.2%** | **11.6%** | **11.3%** | **3.7%** | **7.8%** | **9.0%** | **6.3%** |

**Figure 5.**
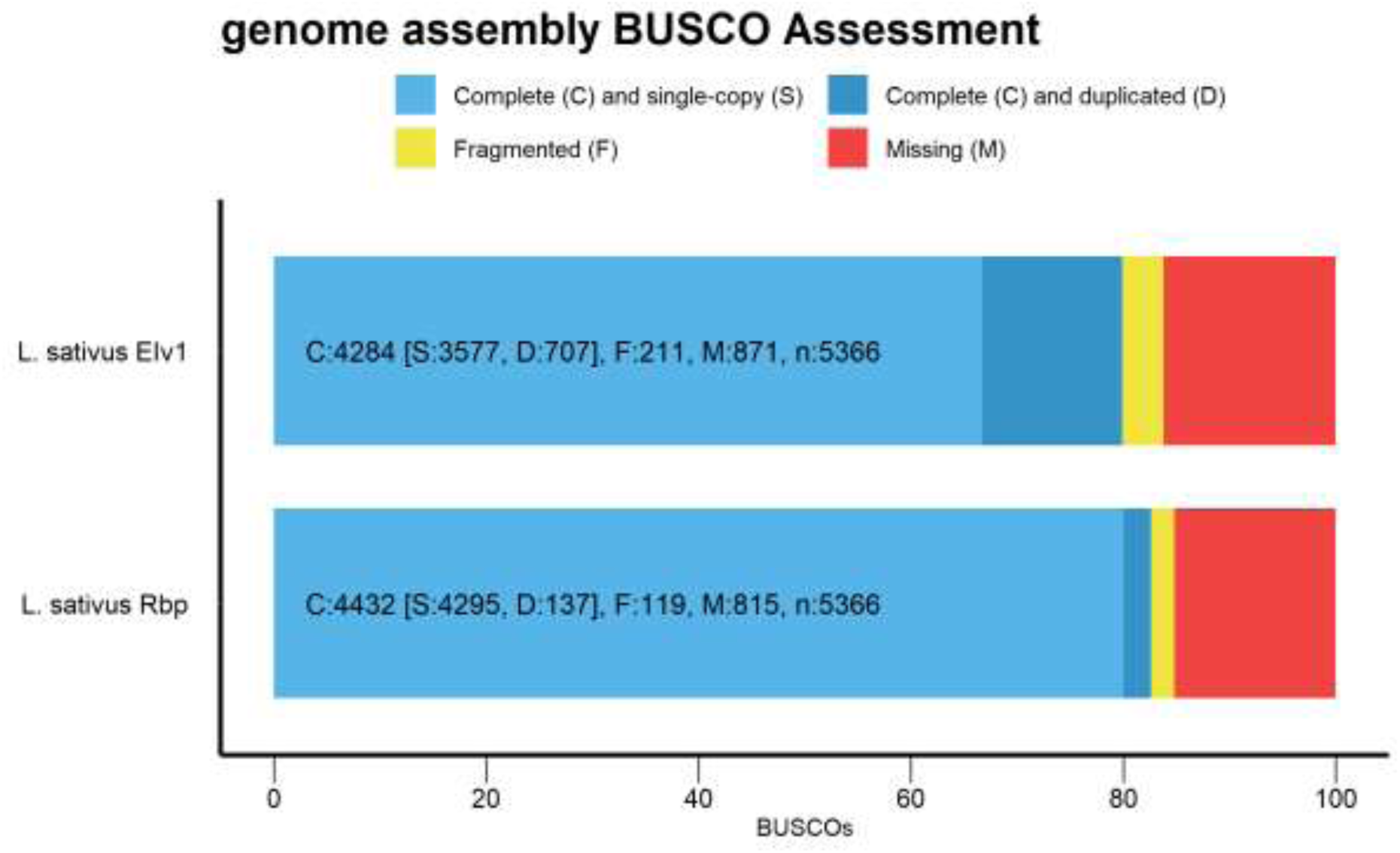
BUSCO (v.4.0.4) assessment of the Lathyrus sativus LS007 genome EIv1 and Rbp assemblies against the Fabales dataset. 5366 total BUSCO groups were searched.

A phylogeny of grass pea (LS007) was determined in the context of 17 other plant species, based on the sequences of 10 BUSCO genes from the viridiplantae set (Figure 6). The sequences for these arbitrary genes are highly similar in both our assemblies causing them to be grouped together in the phylogeny.

**Figure 6.**
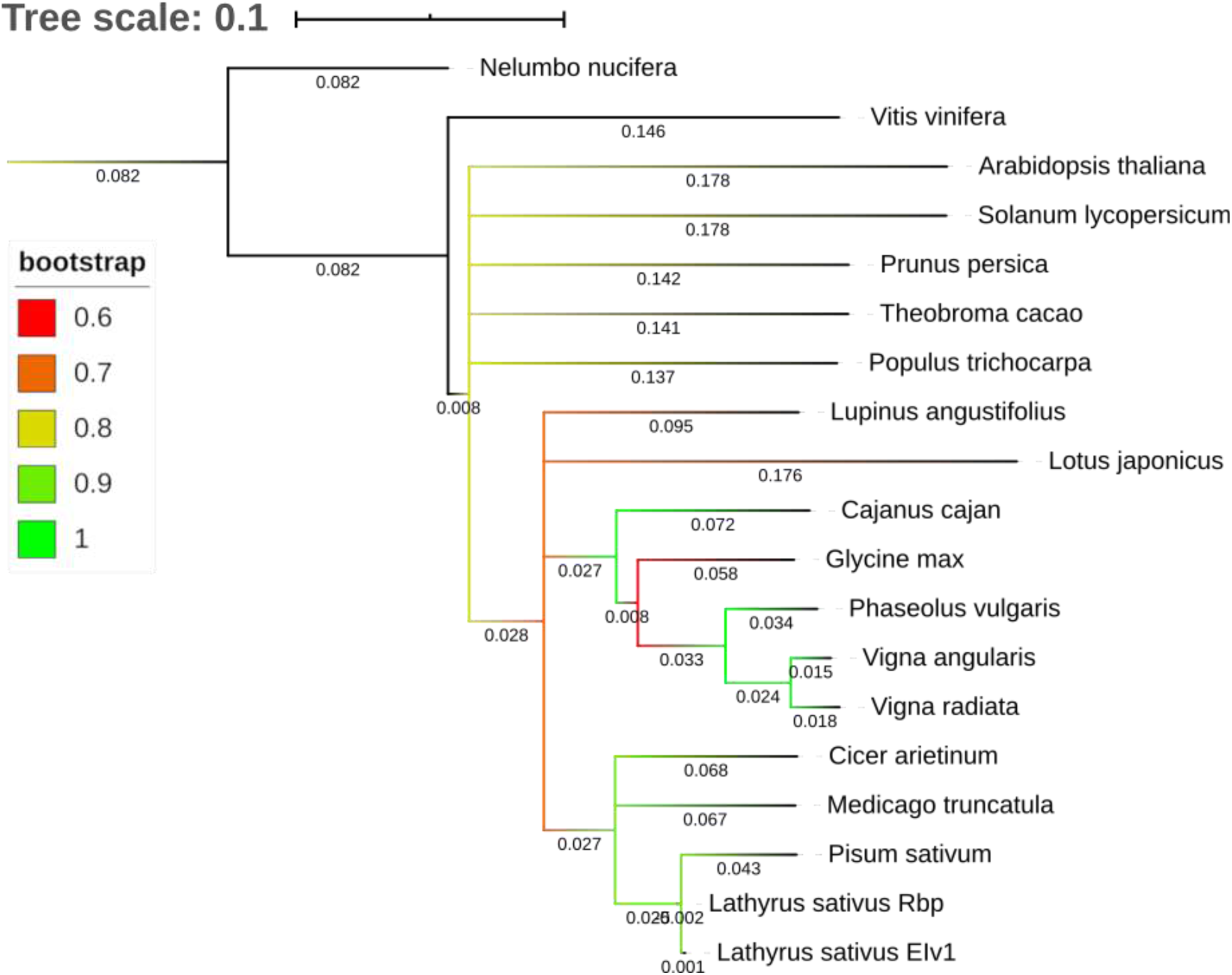
Phylogeny of both LS007 assemblies and 17 other plant species, based on 10 arbitrary BUSCO gene models from the viridiplantae set, re-rooted to Nelumbo nucifera. Colours represent branch bootstrap support. Plotted using iTOL (Letunic and Bork 2019).

### Repeat structure

We screened a subset (0.1 x coverage) of Illumina paired-end reads for repeated elements using the RepeatExplorer2 algorithm. Table 4 shows the portion of reads classified as repeats, along with literature values reported for *Lathyrus sativus* by Macas et al. (2015). As this analysis was done on the basis of the raw read data, it was independent of the assembly strategy used.

**Table 4.**
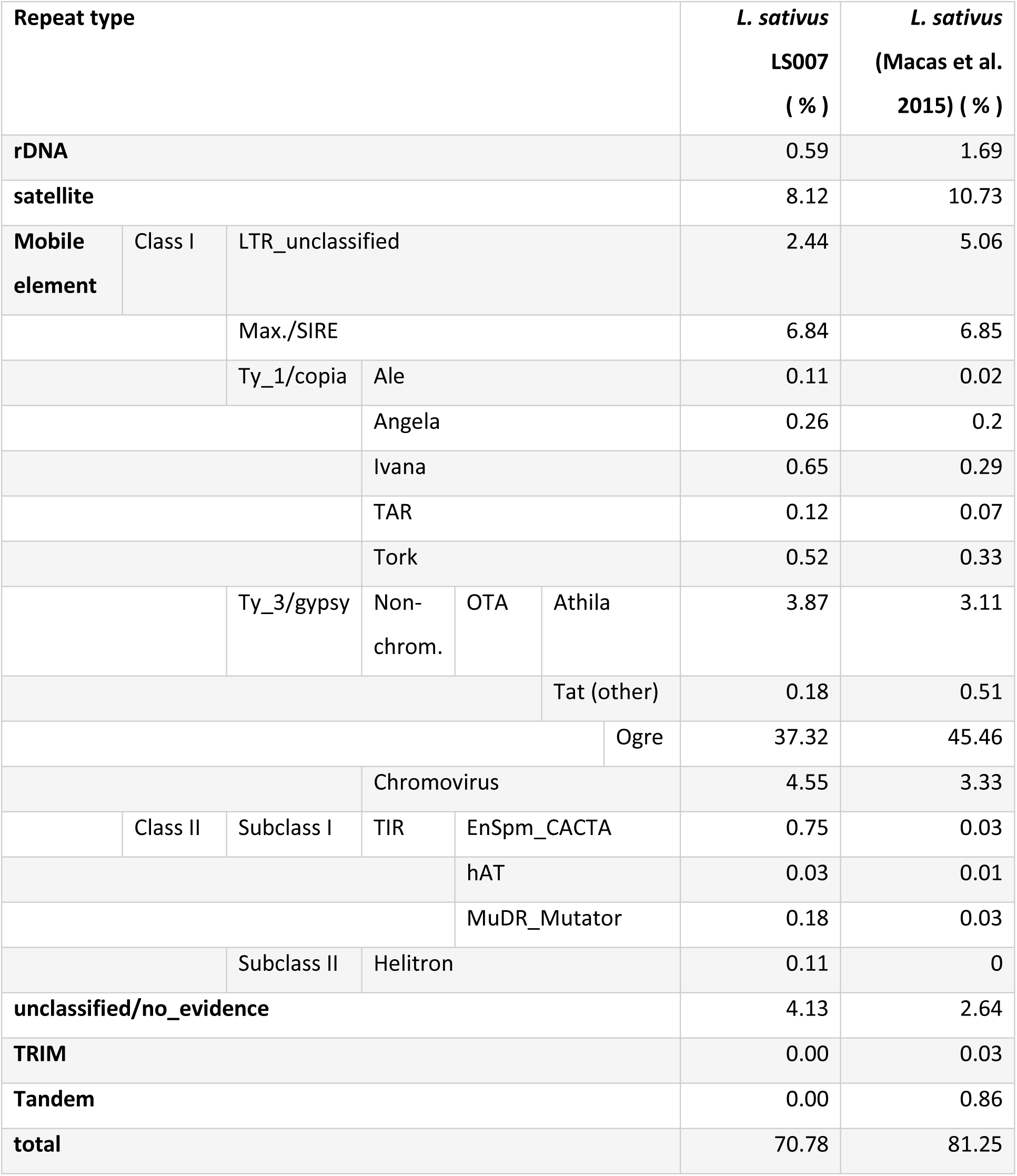
RepeatExplorer2 results summary.

### Annotation

#### EIv1 assembly

We identified a total of 33,819 high confidence gene models in the EIv1 assembly of which 940 were associated with repeats. On average, there were 1.05 transcripts associated with each gene, with a mean cDNA size of 1,454.22 bp and mean transcript size (including introns) of 3,488.6 bp. There were, on average, 4.86 exons per transcript, with an average exon size of 299.17 bp.

A further 53,403 low confidence gene models were also identified, of which 9,577 were associated with repeats. On average, these had 1.03 transcripts associated with each gene, with a mean cDNA size of 752.91 bp and mean transcript size (including introns) of 2645.52 bp. They had, on average, 2.47 exons per transcript, with an average exon size of 305.03 bp.

The number of genes within each category is shown in Table 5. The genome annotation summary statistics are given in Supplemental table S1.

**Table 5.** Annotation biotype and gene confidence assignment for EIv1 assembly.

| Confidence level | Biotype | Gene count |
| --- | --- | --- |
| High | protein_coding | 32,879 |
| High | protein_coding_repeat_associated | 940 |
| Low | protein_coding | 43,826 |
| Low | protein_coding_repeat_associated | 9,577 |

#### tRNA gene prediction

tRNA gene prediction was carried out using tRNAscan-SE (Lowe and Eddy 1997), and resulted in a total of 5765 tRNA genes predicted for the EIv1 assembly and 2801 tRNAs genes predicted for the polished Redbean (Rbp) assembly. Supplemental Table S2 reports the final number of coding transcripts per each rank.

## Discussion

The grass pea genome follows the recent publication of the reference genome of *Pisum sativum* (Kreplak et al. 2019), a close relative of grass pea. Assembling the genome of grass pea is rendered more difficult due to its large size (we estimate 6.3 Gbp for cultivar LS007) and the comparative lack of genetic resources such as genetic and/or molecular maps that could be used for high-level scaffolding to the pseudochromosome level.

Our estimate for the genome size of grass pea (6.3 Gbp) measured by flow cytometry contrasts with the range of values quoted for *Lathyrus sativus* genome size in the literature, including the value listed in the Kew Plant C-Value database of 8.2 Gbp and the ranges given by Ghasem et al. (6.75 Gbp to 7.63 Gbp, (2011)), Ochatt et al. (7.82 to 8.90 Gbp, (2013)), Nandini et al. (6.85 Gbp, (1997)) and Macas et al. (6.52 Gbp (2015)). Legume genomes, particularly species in the Vicaea (=Fabeae) including the genus Lathyrus, are highly variable in size, and this variation correlates strongly with the copy numbers of repeated elements (Leitch et al. 2019; Vondrak et al. 2020). The reported variation in genome size reported for *Lathyrus sativus* could be due to intra-specific variation between genotypes or experimental error. The range of variation in genome size for *Pisum sativum* in the Kew database is 3.49 to 5.42, which is a greater range than for grass pea, and an even greater proportional range (Leitch et al. 2019). As our genome size estimate for LS007 depends on the size of the *Pisum sativum* genome which we used as a standard, the relation between our estimate and the sizes of the assemblies should not be over-interpreted.

Ogre elements are a type of Ty3/gypsy LTR retrotransposon first discovered in legumes (Neumann, Požárková, and Macas 2003; Neumann et al. 2006), but since found in other plant families as well (Macas and Neumann 2007). Ogre elements are characterised by their large size (up to 25 kbp) and the presence of an additional ORF upstream of the gag-pro-pol polyprotein ORF usually present in LTR retrotransposons (Macas and Neumann 2007). In a survey of the genomes of 23 species within the legume family Viciaea (=Fabeae), Ogre elements typically make up about 40% of the entire genome (22.5 - 64.7%), and Ogre-content correlates strongly with genome size (Macas et al. 2015). Our sequencing data gave an estimated Ogre-content of 37% for the LS007 nuclear genome. The repeat analysis using RepeatExplorer2 (Macas et al. 2015) used by both ourselves and Macas et al. relies on <1X coverage raw short-read data and is therefore unaffected by the difficulties of assembling repeat regions that often lead to repeat regions being under-represented in genome assemblies. Nevertheless, our result of 37.3% Ogre-content in LS007 contrasts with the 45.5% estimate reported for *Lathyrus sativus* by Macas et al. (2015). This may be due to genotypic differences between LS007 and the commercial line used by Macas et al.

Satellites in *Lathyrus sativus* have been claimed to originate from the LTRs of Ogre elements (Vondrak et al. 2020). Rapid tandem duplication of such repeated elements may be a mechanism for the high degree of species specificity of satellite elements in legumes. A single satellite element (FabTR53_LAS_A) with a consensus sequence of 660 bp and no significant blastn hits against the NCBI_nt database represented 4.7% of our short reads, corresponding to an estimated 289.8 Mbp or 440,000 copies across the genome. One assembly scaffold carried 144 copies of this satellite in tandem, spread over a 88kbp region. Fluorescence in-situ hybridisation (FISH) conducted by Vondrak et al. (2020) showed that this satellite is concentrated in the sub-telomeric regions of the chromosomes, rather than the primary constrictions of chromosomes, as is typical for most satellites. This high level of repetition underlines the difficulty of assembling the grass pea genome and the necessity for long sequence reads that can span repetitive regions.

Consequently, the assembly of the grass pea genome from short paired-end read data alone reached only a contig N50 of 5.7 kbp, as many repeated regions could not be adequately resolved. Scaffolding with Illumina long mate pair data derived from 2 kbp, 5 kbp, 8 kbp and 14 kbp libraries was able to link contigs into scaffolds reaching an N50 of 59.7 kbp. In the process of scaffolding, many sections of unknown sequence originating from LMP inserts were introduced into the scaffolds, resulting in a total of 1.9 billion Ns in the Eiv1 assembly. It is possible that some smaller contigs may be nested within the unknown sequence of these scaffolds, resulting in a true assembly shorter than the total of 8.12 Gbp. Technical limitations prevent the generation of substantially longer mate pair inserts with reliable sizes. Improving this assembly thus required the use of alternative sequencing and/or scaffolding technologies.

To overcome the challenges of size and repetitiveness of the grass pea genome, we opted for a hybrid assembly approach, combining the high yields and base accuracy of Illumina sequencing with the long reads of Oxford Nanopore sequencing technologies, an approach recommended by Kreplak et al. (2019). Long-read sequencing allowed us to increase assembly contiguity compared to the long-mate-pair-scaffolded Illumina paired-end assembly (155 kbp contig N50 vs. 59 kbp scaffold N50). In addition to the Redbean assembly we report here, we ran Flye (Kolmogorov et al. 2019), which produced a 14 Gbp assembly (before crashing during polishing), and Shasta (Shafin et al. 2019), which produced only a 169 Mbp assembly (results not shown), using the same PromethION data. Neither of these were pursued further. We also ran MaSuRCA (Zimin et al. 2013) using both PromethION and Illumina paired-end data, but the algorithm was unable to complete due to the size of the input datasets.

BUSCO completeness scores were consistently higher in the polished Redbean assembly across 4 BUSCO databases (Simão et al. 2015), although none reached >90% completeness. The polished Redbean assembly exhibited significantly lower rates of duplicated BUSCOs than the EIv1 assembly, on average 4.2% vs. 14.6%. This might reflect duplication of pseudo-hemizygous regions caused by heterozygosity in the sequenced genotype. Although the genotype we used underwent single-seed descent, it is possible that some heterozygotic regions remained in the sequenced material. The longer PromethION reads used by the Redbean assembler collapsed more of these heterozygous regions into single contigs, suggesting that these regions were indeed heterozygous. Duplicated pseudo-hemizygous regions can also be further collapsed using the Purge_dups tool (Guan et al. 2020), but we elected not to do this due to the risk of collapsing truly duplicated genomic regions.

The annotation of the polished Redbean assembly using the transcriptome datasets already described is pending and this draft will be amended accordingly, as soon as it is ready.

The assembly and annotation datasets are available to researchers on request on receipt of a written commitment not to publish analyses of the data before publication of the Consortium paper or on the basis of agreement with all the authors.

## Materials and Methods

### Genome size estimation by flow cytometry

Grass pea genome size was estimated following the procedure described by Dolezel et al. (2007). Fresh, young leaf tissue (40 mg) of grass pea (LS007) and *Pisum sativum* (semi-leafless, obtained from a local market in Nairobi) was sliced finely using a scalpel blade while immersed in 2 mL of ice-cold Galbraith buffer (45 mM MgCl_2_, 30 mM sodium citrate, 20 mM MOPS, 0.1% w/v Triton X-100, pH 7). Three biological replicates were prepared for each grass pea and pea. Supernatants were filtered through one layer of Miracloth (pore size 22-25 µm). One aliquot of 600 µL was prepared from each replicate, along with three grass pea + pea mixes at 2:1, 1:1 and 1:2 ratios, respectively. Propidium iodide was added to each tube to a concentration of 50 µM. Reactions were incubated for 1 h on ice before measuring on a FACSCantoll flow cytometer (Becton Dickinson) with flow rates adjusted to 20-50 events/s. Results were analysed using FCSalyser (v. 0.9.18 alpha). Grass pea genome size was estimated from the mixed sample by dividing the mean of the PE-A peak corresponding to grass pea nuclei by the mean of the PE-A peak corresponding to pea nuclei and multiplying by 4.300 Gbp, the estimated genome size of pea (Leitch et al. 2019).

### Illumina Sequencing

#### Genomic DNA isolation

Seeds of grass pea (*Lathyrus sativus*) were obtained from King’s of Coggershall and underwent six generations of single-seed-descent at the John Innes Centre Germplasm Research Unit. This accession, named LS007 is of European origin, white-flowered, with fully cream-coloured, large and flattened seeds. LS007 seeds are available from the JIC GRU. Seeds were surface sterilised and germinated on distilled water agar in the dark. Genomic DNA was isolated from the etiolated seedlings using a modified CTAB protocol and subsequently, high molecular weight DNA purified using the Qiagen MagAttract kit.

#### Initial Paired end (PE) library DNA sequencing and assembly

Paired end (PE) sequencing was carried out using PCR-free libraries on the HiSeq Illumina platform. Five lanes of untrimmed PE data from PSEQ-907 (LIB21060) were assembled using both Abyss (Simpson et al. 2009) and Discovar de novo v.52488 (Love et al. 2016). A range of kmer spectra plots were generated using KAT (Mapleson et al. 2017) to determine how the kmers from the initial PE library were represented in these final PE assemblies. Given broad similarity between kmer spectra for both assemblies, and that the best assembly statistics were returned by Discovar de novo (best Abyss N50 = 1.5 kbp, best Discovar de novo N50 = 5.7 kbp), the Discovar de novo assembly was selected as the best candidate for further improvement/scaffolding. The total usable coverage from clipped LMP libraries (using categories A, B and C as identified by *Nextclip*) are given in Supplemental table S3

#### Sequencing and assembly of Long Mate Pair (LMP) libraries

A pool of Long Mate Pair (LMP) libraries with a range of insert sizes (12 in total) was prepared and sequenced. These reads were processed with *nextclip*, and mapped to both the best Abyss and Discovar de novo assemblies. Binary Alignment Map (BAM) files were sorted and de-duplicated before running the Picard tool *CollectInsertSizeMetrics* on them (Wysokar et al. 2016). The Discovar de novo contigs/scaffolds were generally longer and therefore judged to be the better set for mapping, particularly for the LMPs with longer insert sizes. The insert size estimates from *CollectInsertSizeMetrics* were supplemented with additional statistics from the Picard *JumpingLibraryMetrics* module. Analysis of both these sets of metrics suggested that the LIB21840 (∼2 kbp), LIB21836 (∼5 kbp) and LIB21834 (∼8 kbp) LMPs were the three best libraries for resequencing for greater coverage depth.

These three LMP libraries were pooled (IPO3823) and sequenced in four separate lanes across three different sequencing runs (PSEQ-1057, PSEQ-1067 and PSEQ-1101). *Nextclip* was used on the reads after each run to assess the extent to which usable LMP coverage had been recovered.

#### Scaffolding

After an initial scaffolding attempt using SSPACE v2.0 (Boetzer et al. 2011), an extra lane of the longest 8 kbp LMP (LIB21834) was sequenced on PSEQ-1159, and this helped to increase the final clipped coverage for this library to 10.7x (assuming a genome size of 6.3 Gbp). Use of the Bowtie alignment option of SSPACE (v3.0) was rapidly judged to be the only feasible option (because Bwa was extremely slow), but this necessitated making a custom installation of SSPACE which was able to handle genome references of >4 Gbp. This version of SSPACE was run specifying k=3 as the minimum number of connections required before scaffolding two sequences together. All contigs/scaffolds of >500 bp from the Discovar de novo assembly were provided as the input reference. The final scaffold N50 using 500 bp as the input length cutoff was ∼32 kbp. This assembly was re-scaffolded using an even longer insert LMP (LIB28873, ∼14 kbp insert size). Using exactly the same scaffolding procedure and parameters, providing only the clipped LIB28873 reads (final coverage is 5.2x assuming 6.3 Gbp genome size), the final scaffold N50 was ∼47 kbp. Increasing the minimum length to 1 kbp improved the assembly, resulting in a final draft assembly of 8.12 Gbp (comprising 6.2 Gbp known nucleotides and 1.9 Gbp Ns) and a scaffold N50 of 59.7 kbp. The final assembly statistics are shown in Table 2 and Figure 4.

Annotation of the Illumina assembly (EIv1)The gene annotation pipeline used to annotate the LS007 EIv1 genome assembly has been adapted from the pipeline used to annotate the wheat genome developed at the Earlham Institute (Venturini, Kaithakottil, and Swarbreck 2016).

#### Identification of repeats from the genome assembly

RepeatModeler (v1.0.10 - http://www.repeatmasker.org/RepeatModeler/) was used for *de novo* identification of repetitive elements from the assembled grass pea genome sequence. Protein coding genes in the RepeatModeler generated library were hard-masked (i.e. replaced with Ns) using the Arabidopsis Araport11 dataset and *Cicer arietinum* Annotation 101 (https://www.ncbi.nlm.nih.gov/genome/annotation_euk/Cicer_arietinum/101/) coding genes. Any genes with descriptions indicating “transposon” or “helicase” were removed. TransposonPSI (r08222010) (Haas 2010) was run and significant hits hard-masked, with the output being used to mask the RepeatModeler library. Unclassified repeats were searched in a custom BLAST database of organellar genomes (mitochondrial and chloroplast sequences from Fabales [ORGN] in NCBI nucleotide division, downloaded on 22.9.2017). Any repeat families matching organellar DNA were also hard-masked.

Repeat identification was refined by running RepeatMasker v4.0.7 (Smit, Hubley, and Green 2013) with RepBase Viridiplantae library and with the customized repeatmodeler library (i.e. after masking out protein coding genes), both using the -nolow setting.

The combined masking resulted in ∼62% of the grass pea genome being soft-masked (i.e. rendered in lowercase to stop alignment of transcriptome data).

### Reference guided transcriptome reconstruction

#### Alignment of Illumina RNA-Seq data

RNA-Seq data from three different genotypes LS007, LSWT11 and Mahateora (Emmrich 2017; Emmrich et al. 2019) was used for grass pea genome annotation: 12 samples from LS007 and Mahateora each (each root and shoot tissues from 3 biological replicates of droughted and non-droughted treatments) and 7 samples from seedling shoot tip, seedling root tip, whole root, whole leaves, early flowers, early pods and late pods from LSWT11 (Table 2) for a total of 31 individual RNA-seq samples). In total, the three filtered datasets comprised over 2.5 billion paired-end reads. For each dataset, read samples were collapsed by tissue and filtered using Xtool (Nazar et al. 2020), with the command line options. Due to concerns about high concentrations of ribosomal RNA, datasets were further filtered using SortMeRNA v. 2.0 (Kopylova, Noé, and Touzet 2012), and using RFam (5S and 5.8S) and Silva (Archaea 16S-23S, Bacteria 16S-23S, Eukaryota 18S-28S) as databases.

#### Alignment with HISAT2

Filtered reads were aligned to the grass pea genome using HISAT2 v2.0.5 (Kim, Langmead, and Salzberg 2015, 2) with options: --min-intronlen=20 --max-intronlen=50000 --dta-cufflinks. The RNA-seq data and alignments are summarised in supplemental table S4.

#### Transcript assembly

The Illumina RNA-Seq alignments (31 RNA-Seq transcript alignments from HISAT2) were assembled using two different RNA-Seq assembly tools: Cufflinks v. 2.2.1 (Trapnell et al. 2010; Roberts et al. 2011), with options: --min-intron-length=20 --max-intron-length=50000 and StringTie v.1.3.3 (Pertea et al. 2015), with options: -f 0.05 -m 200. The results of transcript assembly are shown in supplemental table S5.

Mikado v1.0.1 (Venturini et al. 2018), which uses transcript assemblies generated by multiple methods to improve transcript reconstruction, was used to integrate the ∼4.5 million Illumina contigs generated using Cufflinks and StringTie (see supplemental table S6). Loci were first defined across all assemblies, followed by scoring of each transcript for ORF and cDNA length, position of the ORF within the transcript (and presence of multiple ORFs), UTR lengths. The highest scoring transcript assembly was used along with any additional transcripts (splice variants) compatible with this representative transcript assembly. Mikado selected transcript sets were generated for use in gene predictor training and annotation (Table 4) using the RNA-Seq transcript assemblies with the “chimera_split” option set to PERMISSIVE. For Mikado runs incorporating BLAST data, transcripts that had passed the “prepare” step were compared to filtered and masked proteins from *Cicer arietinum, Cucumis sativus, Fragaria vesca, Glycine max, Malus domestica, Medicago truncatula, Prunus persica, Phaseolus vulgaris, Trifolium pratense* using BLAST+ v. 2.6.0 (Altschul et al. 1997). Each result was limited to the best 15 matches.

#### Gene prediction using evidence guided AUGUSTUS

Protein coding genes were predicted using AUGUSTUS (Stanke et al. 2004; 2006), which uses a Generalized Hidden Markov Model (GHMM) employing both intrinsic and extrinsic information.

##### Intron/exon junctions

RNA-Seq junctions were derived from RNA-Seq alignments using Portcullis v1.0.2 (Mapleson et al. 2018) with default filtering parameters. Junctions that passed the Portcullis filter with a score of 1 were classified as ‘Gold’ and with a score <1 were classified as ‘Silver’.

##### Proteins

Predicted protein sequences from 9 species (*Cicer arietinum, Cucumis sativus, Fragaria vesca, Glycine max, Malus domestica, Medicago truncatula, Prunus persica, Phaseolus vulgaris, Trifolium pratense*) were soft-masked for low complexity regions (segmasker from NCBI BLAST+ 2.6.0) and aligned to the soft-masked grass pea genome sequence (using repeatmodeler repeats) with exonerate v2.2.0 (Slater and Birney 2005) with parameters: --model protein2genome -- showtargetgff yes --showvulgar yes --softmaskquery yes --softmasktarget yes --bestn 10 --minintron 20 --maxintron 50000 --score 50 --geneseed 50.

To identify high confidence alignments, exonerate results were filtered at 50% identity and 80% coverage. Alignments with introns longer than 10 kbp were removed from further analyses (see Supplemental table S7).

##### Classification of Mikado transcripts

The primary Mikado models for each locus were classified into three categories:

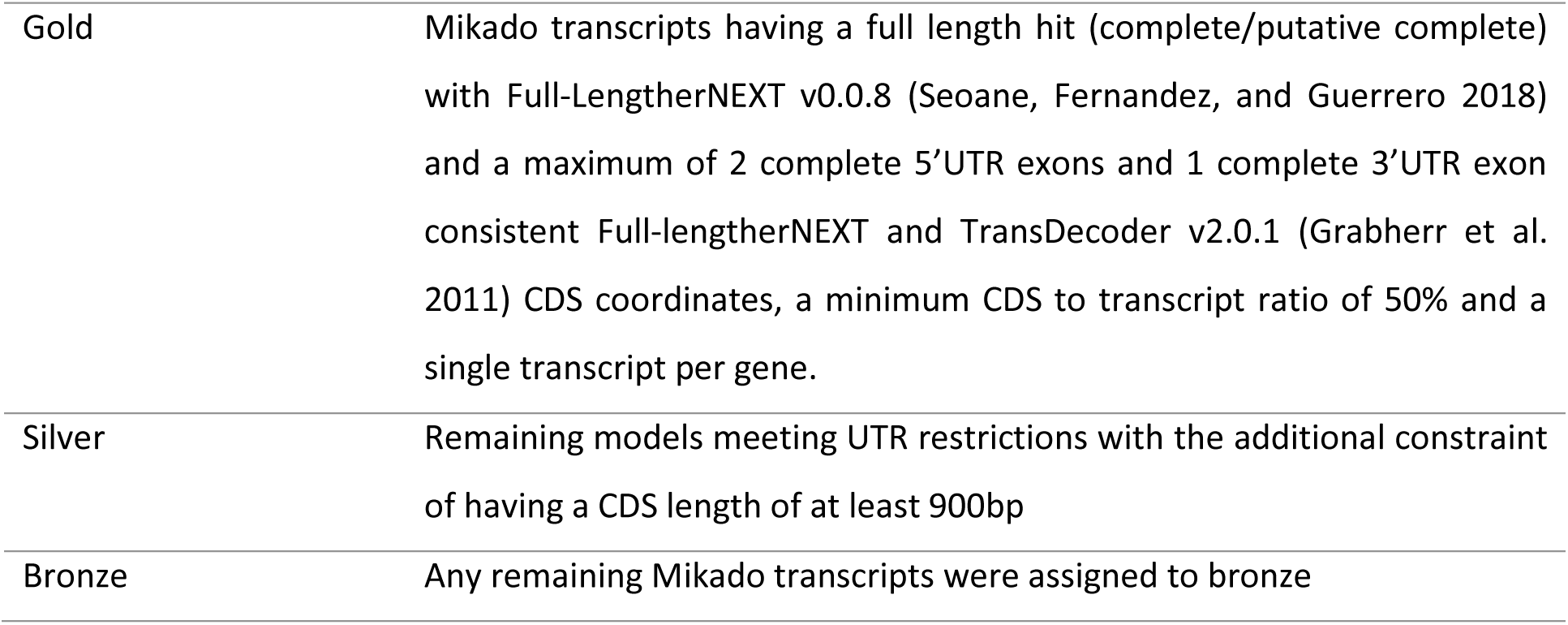

### Gene prediction

#### Gene predictor training

The Mikado gold set transcripts were selected for training AUGUSTUS (Grabherr et al. 2011).We excluded genes with a genomic overlap within 1000bp of a second gene and gene models that were homologous to each other with coverage and identity ≥ 80%. The filtered set contained 9604 transcripts, from which 2000 transcripts were selected at random for training AUGUSTUS and another 200 transcripts were used for testing. The trained AUGUSTUS model resulted in values of 0.976 sensitivity (sn), 0.909 specificity (sp) at the nucleotide level, sn 0.837, sp 0.801 at the exon level and sn 0.41, sp 0.376 at the gene level.

AUGUSTUS (v2.7) was used to predict gene models for the EIv1 assembly using the evidence hints generated from nine sets of cross species protein alignments (listed above), Mikado Illumina models and intron/exon junctions defined using the RNA-Seq data. Interspersed repeats were provided as “nonexonpart” to exclude them from analysis. We assigned additional bonus scores and priority based on evidence type and classification (Gold, Silver, Bronze) to reflect the reliability of different evidence sets.

#### Cross species protein similarity ranking

Each gene model was assigned a protein rank (P1–P5) reflecting the level of coverage of the best identified homolog in plant protein databases. Protein ranks were assigned according to:

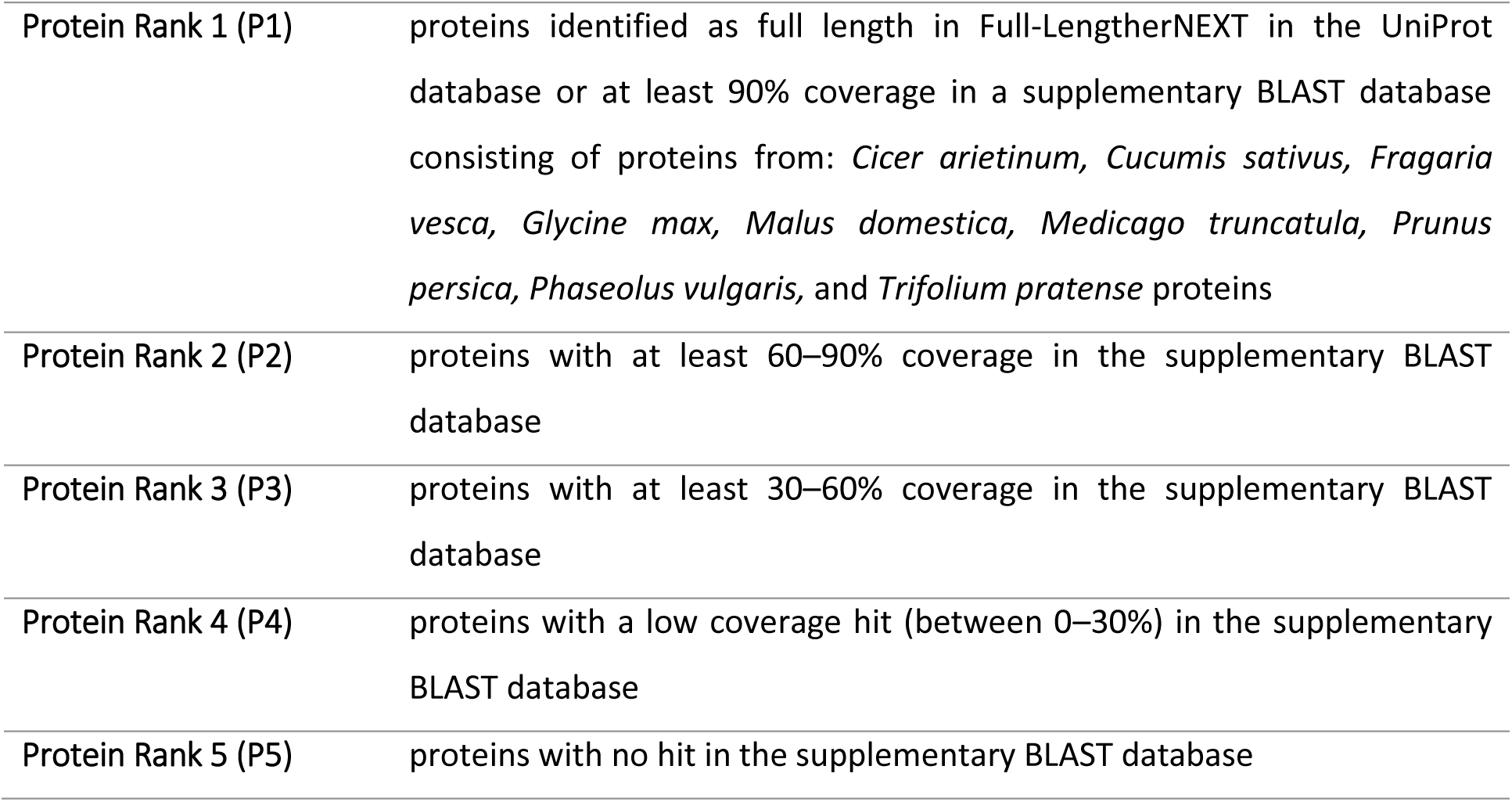

|  |  |
| --- | --- |
| Protein Rank 1 (P1) | proteins identified as full length in Full-LengtherNEXT in the UniProt database or at least 90% coverage in a supplementary BLAST database consisting of proteins from: *Cicer arietinum*, *Cucumis sativus*, *Fragaria vesca*, *Glycine max*, *Malus domestica*, *Medicago truncatula*, *Prunus persica*, *Phaseolus vulgaris*, and *Trifolium pratense* proteins |
| Protein Rank 2 (P2) | proteins with at least 60–90% coverage in the supplementary BLAST database |
| Protein Rank 3 (P3) | proteins with at least 30–60% coverage in the supplementary BLAST database |
| Protein Rank 4 (P4) | proteins with a low coverage hit (between 0–30%) in the supplementary BLAST database |
| Protein Rank 5 (P5) | proteins with no hit in the supplementary BLAST database |

#### Grass pea transcript support ranking

Transcript ranks (T1–T5) were assigned based on the support for each predicted gene model from the assembled grass pea RNA-Seq data (all 4,582,819 transcripts assembled using Cufflinks and StringTie).

We calculated a variant of annotation edit distance (*AED*) and used this to determine a transcript level ranking.

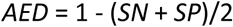

where *SN* is sensitivity and *SP* specificity.

*AED* was calculated at base, exon and splice junction levels against all individual transcripts used in our gene build (Illumina assemblies). The mean of base, exon and junction *AEDs* (derived using the Mikado ‘compare utility’) based on the transcript that best supported the gene model was then used to assign transcript rank as shown:

|  |  |
| --- | --- |
| **Transcript Rank 1 (T1)** | Full length support single cDNA reads (not relevant to this study, but retained for workflow compatibility) |
| **Transcript Rank 2 (T2)** | full length support from Illumina assemblies |
| **Transcript Rank 3 (T3)** | Best average *AED* less than 0.5 |
| **Transcript Rank 4 (T4)** | Best average *AED* between 0.5 and 1 |
| **Transcript Rank 5 (T5)** | No transcriptomic support (best average *AED* = 1) |

#### Assignment of gene biotypes and confidence classification

Gene models were classified as either coding or repeat associated and classified as high- or low-confidence based on cross-species protein similarity and grass pea transcript evidence. Each transcript was assigned a ‘protein rank’ and a ‘transcript rank’ (as described above) and a binary confidence tag (“High” vs “Low”), combining the two. Any transcript with protein rank P2 or better as well as transcript rank T4 or better were classified as ‘high confidence’, all others as ‘low confidence’.

#### Assignment of a locus biotype

Using the protein and transcript rankings, we assigned a locus biotype (‘repeat-associated’ or ‘protein-coding’) to each gene.

##### Repeat-associated biotypes

Genes were classified as ‘repeat-associated’ if ANY of their transcripts: :

- aligned with at least 20% similarity and 30% coverage to the TransposonPSI library v08222010 (Haas 2010)
- or had at least 30% coverage by RepeatModeler/RepeatMasker derived interspersed repeats
- or had a match for “retrotransposon”, “transposon”, “helicase” in their annotation (AHRD and interproscan) description.

##### Protein-coding genes

Genes not assigned as ‘repeat-associated’, were assigned the ‘protein coding’ biotype, along with all transcripts associated with them.

#### Removal of spurious genes

After assigning biotypes to each gene, we used Kallisto v 0.43.1 (Bray et al. 2016) to estimate gene expression in all of our samples. Genes were retained if one of their transcripts had either an expression level exceeding 0.5 transcripts per million (TPM) in at least one sample, as measured by Kallisto OR any BLAST hits from the Full-LengtherNEXT analysis against the UniProt database.

Any gene whose transcripts were all marked for removal was excluded from the final annotation.

#### Assignment of a representative gene model

We assigned representative gene models by selecting the model with the highest confidence ranking (as shown in Supplemental table S2), and the lowest *AED* with the order of priority for ranking:

1. highest protein rank
2. highest transcript rank
3. lowest *AED*.

#### Functional annotation of protein coding transcripts

Proteins were annotated using AHRD v.3.1 (Hallab 2014). Predicted protein sequences were searched against Araport11 *A. thaliana* protein sequences (Cheng et al. 2017) and the plant sequences of UniProt v. 23Nov2017, both SwissProt and TREMBL datasets (The UniProt Consortium 2019) using BLASTP+ v. 2.6.0 asking for a maximum e-value of 1e-5. We also ran InterProScan 5.22.61 (Jones et al. 2014) and provided the InterProScan output to AHRD. We adapted the standard example configuration file pathtest/resources/ahrd_example_input.yml, distributed with the AHRD tool by providing the GOA mapping from UniProt. Supplemental table S2 reports the final number of coding transcripts per each rank.

### PromethION Sequencing

Two methods of DNA extraction and three methods of DNA processing were used to optimise yield and read length profile of the PromethION sequencing.

#### DNA extraction

##### Circulomics Nanobind Plant Nuclei Kit extraction (CN)

Fresh, aseptic LS007 shoot tissue (0.5 g) was ground in liquid nitrogen using a mortar and pestle. The resulting powder was resuspended by vortexing in 10 mL of nuclei extraction buffer (0.35 M Sorbitol, 100 mM Tris pH 8.0, 5mM EDTA and 1% β-mercaptoethanol) and then filtered through Miracloth. The filtrate was centrifuged at 750 x g for 15 min at 4 °C and the pellet was resuspended in nuclei extraction buffer (with the addition of 0.4% Triton). The wash was repeated before centrifugation and resuspension in 1mL of nuclei resuspension buffer (0.35 M Sorbitol, 100 mM Tris, 5mM EDTA). The suspension was then centrifuged at 750 x g for 15 min at 4 °C and the supernatant was removed. The resulting nuclear pellet was taken into the Circulomics Nanobind Plant nuclei Kit (Circulomics; SKU NB-900-801-01) according to the manufacturers protocol. The resulting DNA had a peak fragment size >60 kbp.

##### Qiagen DNeasy Plant Mini Kit extraction (QD)

0.5 g of fresh, aseptic LS007 shoot tissue was ground under liquid nitrogen using a mortar and pestle and resuspended in 2 mL of AP1 (Qiagen) with 20 µl of RNase I (Qiagen). This was incubated at 65°C for 10 minutes, before aliquoting into 5 tubes and proceeding with the Qiagen DNeasy Plant Mini Kit (Qiagen; 69104) protocol. Eluted DNA was pooled into a single tube and had a peak fragment size of 48 kbp.

#### DNA processing

The process of Sample preparation was iteratively optimized to achieve both high total sequence yield and long read length. The exact combination of techniques used for each library preparation is given in Table 1.

##### Ampure XP bead purification/concentration (AMP)

To increase the concentration of DNA for input into the Short Read Eliminator and to reduce the loss of DNA in library preparation, All except the DNA Qiagen DNeasy-extracted samples were pre-incubated with 0.8x Ampure XP (Beckman Coulter; A63881). Beads were washed twice in 80% ethanol and DNA was eluted from beads with TE buffer (10 mM Tris HCl pH 8.0, 1 mM EDTA).

##### Needle shearing (NS)

Starting with the 2^nd^ loading of flowcell FC1, all except the Qiagen DNeasy-extracted samples were needle-sheared 30 times with a 26 gauge needle to reduce the amount of very high molecular weight DNA, which can cause blocking and therefore an artificially low N50.

##### Short Read Elimination (SRE)

Starting with the 3^rd^ loading of flowcell FC1, all samples were subjected to a Circulomics Short Read Eliminator (Circulomics; SKU SS-100-101-01) treatment to reduce the number of short fragments.

#### Library preparation and loading

All libraries were prepared using the Genomic DNA by Ligation (SQK-LSK109) – PromethION Kit (Oxford Nanopore Technologies; SQK-LSK109) following the manufacturer’s procedure. Libraries were loaded onto PromethION Flow Cells (Oxford Nanopore Technologies; FLO-PRO002) at between 250-400 ng. Due to the rapid accumulation of blocked flow cell pores or due to apparent read length anomalies on some runs, flow cells used in runs were treated with a nuclease flush to digest blocking DNA fragments before reloading with fresh library according the Oxford Nanopore Technologies Nuclease Flush protocol, version NFL_9076_v109_revD_08Oct2018.

### Data processing

Fast5 sequences produced by PromethION sequencing were basecalled using the Guppy high accuracy basecalling model (dna_r9.4.1_450bps_hac.cfg) and the resulting fastq files were quality filtered by the basecaller. Fastq files from all five sequencing runs were pooled and assembled using Redbean (Ruan and Li 2020, 2) (previously wtdbg2, version 2.2). This assembly was polished with minimap2 (v. 2.17) (Li 2018, 2) using the original nanopore dataset in fasta format, followed by polishing using bwa (v. 0.7.17) (Li 2013).

### BUSCO analysis

Gene space completeness in Rbp and EIv1 was assessed using BUSCO v.4.0.4_cv1 (Simão et al. 2015) and the odb10 databases for eukaryta, viridiplantae, eudicots and fabales, employing default parameters.

### Phylogeny

Ten arbitrary single-copy BUSCO sequences found in the Rbp and EIv1 assemblies and their predicted mRNA homologs with the highest NCBI BLAST scores in *Arabidopsis thaliana, Cajanus cajan, Cicer arietinum, Glycine max, Lotus japonicus, Lupinus angustifolius, Medicago truncatula, Nelumbo nucifera, Phaseolus vulgaris, Pisum sativum, Populus trichocarpa, Prunus persica, Solanum lycopersicum, Theobroma cacao, Vigna angularis, Vigna radiata* and *Vitis vinifera* (Sayers et al. 2019). Multiple sequence alignments for each homolog group were produced using MUSCLE (Edgar 2004). We merged the 10 unrooted trees each containing 19 unique taxa (including the two LS007 assemblies, EIv1 and Rbp) with 79 unique splits from total of 360, using DendroPy 3.12.0 (Sukumaran and Holder 2010), and the consensus tree was built using the minimum clade frequency threshold set at 0.5.

### Repeat analysis

Genome repeat structure was analysed using RepeatExplorer2 (Macas et al. 2015), a graph-based read clustering algorithm for repeat identification. Illumina HiSeq paired end data was downsampled to a coverage of 0.1X and trimmed, quality filtered, cutadapt filtered and interlaced into a single FASTA file for processing by repeatExplorer2 using the ELIXIR CZ Galaxy server using default parameters (Afgan et al. 2016).

## Acknowledgments

This work was supported by a John Innes Centre Institute Development Grant, the Biotechnology and Biological Sciences Research Council (BBSRC) Detox Grass pea project (BB/L011719/1) the BBSRC SASSA UPGRADE project (BB/R020604/1), the BBSRC Institute Strategic Programme (BBS/E/J/000PR9799) and the Nottingham Future Food Beacon of Excellence. PMFE’s studentship was funded by the John Innes Foundation’s Student Rotation programme. None of the funding bodies were involved in the design of this study, the collection or analysis or interpretation of data, or in writing the manuscript.

The Galaxy server that was used for some calculations is in part funded by Collaborative Research Centre 992 Medical Epigenetics (DFG grant SFB 992/1 2012) and German Federal Ministry of Education and Research (BMBF grants 031 A538A/A538C RBC, 031L0101B/031L0101C de.NBI-epi, 031L0106 de.STAIR (de.NBI)).

We acknowledge Guru Radhakrishnan, Tjelvar Olsen, Matt Hartley, Shabhonam Caim, Pirita Paajanen and Burkhard Steuernagel for their vital support and advice in bioinformatics and data handling.

## Availability of data and materials

All plant materials used in this article are available from the corresponding author. The datasets used and analysed during the current study have been uploaded (under embargo) to the European Nucleotide Archive but are available from the corresponding author upon reasonable request. All data will become publicly available upon publication of the peer-reviewed article.

