## Supplementary tables 1, 2, 3, 4, 5, 6 and 7 for "A draft genome of grass pea (*Lathyrus sativus*), a resilient diploid legume"

Table S1. AUGUSTUS Gene model summary statistics for Elv1 assembly. Gene models are informed by transcript assemblies (Supplemental table S6) and alignments of reference proteins (Supplemental table S7)

| | All gene models | High confidence gene models | Low confidence gene models |
|---|---|---|---|
| **Gene Count** | 87,222 | 33,819 | 53,403 |
| **Total transcripts** | 90,253 | 35,500 | 54,753 |
| **transcripts per gene** | 1.03 | 1.05 | 1.03 |
| **Transcript mean size (inc. introns) (bp)** | 2,977.14 | 3,488.60 | 2,645.52 |
| **Transcript mean size cDNA (bp)** | 1,028.76 | 1,454.22 | 752.91 |
| **Transcript median size cDNA (bp)** | 818 | 1,249 | 671 |
| **cDNA minimum size (bp)** | 31 | 124 | 31 |
| **cDNA maximum size (bp)** | 18,353 | 18,353 | 9,517 |
| **Total exons** | 307,706 | 172,558 | 135,148 |
| **Exons per transcript** | 3.41 | 4.86 | 2.47 |
| **Exon mean size (bp)** | 301.74 | 299.17 | 305.03 |
| **Total distinct exons** | 294,343 | 163,977 | 130,452 |
| **Distinct exon mean size (bp)** | 306.78 | 304.04 | 310.14 |
| **CDS mean size (bp)** | 236.25 | 232.96 | 240.67 |
| **Distinct CDS mean size (bp)** | 239.31 | 235.98 | 243.68 |
| **Transcript mean CDS size (bp)** | 744.95 | 1,069.94 | 534.23 |
| **Transcript median CDS size (bp)** | 561 | 885 | 452 |
| **CDS minimum size (bp)** | 3 | 9 | 3 |
| **CDS maximum size (bp)** | 18,098 | 18,098 | 9,224 |
| **5' UTR mean size (bp)** | 121.60 | 142.59 | 104.37 |
| **Distinct 5' UTR mean size (bp)** | 121.77 | 142.81 | 104.44 |
| **3' UTR mean size (bp)** | 181.18 | 221.90 | 149.57 |
| **Distinct 3' UTR mean size (bp)** | 181.12 | 222.15 | 149.65 |

Table S2. Rankings and confidence assignments for coding transcripts in the EIv1 assembly.

| Protein Rank | Transcript Rank | Confidence | Gene count | Transcript count |
|---|---|---|---|---|
| P1 | T2 | High | 19,622 | 19,862 |
| P1 | T3 | High | 5,394 | 5,916 |
| P1 | T4 | High | 512 | 556 |
| P1 | T5 | Low | 2,872 | 2,935 |
| P2 | T2 | High | 4,814 | 5,000 |
| P2 | T3 | High | 3,105 | 3,422 |
| P2 | T4 | High | 372 | 390 |
| P2 | T5 | Low | 3,591 | 3,629 |
| P3 | T2 | Low | 5,651 | 5,904 |
| P3 | T3 | Low | 5,752 | 6,161 |
| P3 | T4 | Low | 845 | 882 |
| P3 | T5 | Low | 5,577 | 5,617 |
| P4 | T2 | Low | 4,168 | 4,326 |
| P4 | T3 | Low | 6,211 | 6,529 |
| P4 | T4 | Low | 1,069 | 111 |
| P4 | T5 | Low | 6,400 | 6,428 |
| P5 | T2 | Low | 2,054 | 1,092 |
| P5 | T3 | Low | 3,194 | 3,417 |
| P5 | T4 | Low | 1,210 | 1,256 |
| P5 | T5 | Low | 4,809 | 4,818 |

Table S3. Total sequence coverage from clipped LMP libraries (using categories A, B and C as identified by Nextclip). Values for LIB21834 and LIB28873 are approximate.

| Library | Insert size | Clipped LMP bases | total coverage (6.3 Gbp) |
|---|---|---|---|
| **LIB21840** | 2 kbp | 60,980,674,152 | 9.68 |
| **LIB21836** | 5 kbp | 42,279,763,347 | 6.71 |
| **LIB21834** | 8 kbp | 67,410,000,000 | 10.7 |
| **LIB28873** | 14 kbp | 32,760,000,000 | 5.2 |

Table S4. Total RNA-seq reads for LS007, Mahateora, and LSWT11 and summary of their alignment against the EIv1 LS007 genome assembly using HISAT2.

| | **LS007** | **Mahateora** | **LSWT11** |
|---|---|---|---|
| **Number of samples** | 12 | 12 | 7 |
| **Number of filtered reads** | 619,136,758 | 608,250,254 | 1,299,302,860 |
| **Mean fil. rds. per sample** | 51,594,730 | 50,687,521 | 185,614,694 |
| **Aligned reads (HISAT2)** | 567,207,986 | 551,021,439 | 1,160,184,405 |
| **Aligned reads percentage** | 91.61% | 90.59% | 89.29% |

Table S5. Illumina transcript assembly statistics showing Cufflinks and StringTie assembly results. Assemblies were run using the 31 alignments produced by HISAT2 (one for each sample), shown in Table S4. The transcript number shown represents the total number (redundant set) of transcripts each assembler generated. For each tool, assembled transcripts have been clustered into loci using cuffcompare (cufflinks v2.2.1; command line options "-C -G").

| Method | Loci | Transcripts | Mean number of exons | Mean cDNA size | Number of monoexonic transcripts |
|---|---|---|---|---|---|
| **Cufflinks** | 104,302 | 1,892,101 | 3.70 | 1188.24 | 690,319 |
| **StringTie** | 158,635 | 2,690,718 | 3.40 | 1090.66 | 1,070,394 |

Table S6. Mikado transcript assembly statistics for EIv1 assembly. Mikado unifies data generated by Cufflinks and StringTie for each of the 31 alignments one non-redundant set of transcripts (assmblies are summarized in Table S5).

| Method | Loci | Transcripts | Mean number of exons | Mean cDNA size | Number of monoexonic transcripts |
|---|---|---|---|---|---|
| **Mikado** | 111,340 | 144,721 | 3.29 | 904.95 | 100,071 |

Table S7. Reference protein datasets used with AUGUSTUS. Proteins were filtered at 50% identity and 80% coverage. Any intron over 10 kbp resulted in the protein alignment being removed.

| | ***Cicer arietinum*** | ***Cucumis sativus*** | ***Fragaria vesca*** | ***Glycine max*** | ***Malus domestica*** | ***Medicago truncatula*** | ***Prunus persica*** | ***Phaseolus vulgaris*** | ***Trifolium pratense*** |
|---|---|---|---|---|---|---|---|---|---|
| **Total proteins** | 33,107 | 30,364 | 32,831 | 88,647 | 63,517 | 62,319 | 47,089 | 36,995 | 41,297 |
| **proteins aligned** | 22,890 | 14,575 | 7,399 | 53,077 | 18,209 | 32,739 | 20,221 | 22,653 | 24,122 |
| **Proteins aligned %** | 69.1% | 48.0% | 22.5% | 59.9% | 28.7% | 52.5% | 42.9% | 61.2% | 58.4% |
| **Protein alignments** | 55,133 | 37,114 | 19,767 | 128,682 | 51,884 | 88,027 | 49,910 | 56,122 | 70,681 |
